# R2d2 drives selfish sweeps in the house mouse

**DOI:** 10.1101/024851

**Authors:** John P Didion, Andrew P Morgan, Liran Yadgary, Timothy A Bell, Rachel C McMullan, Lydia Ortiz de Solorzano, Janice Britton-Davidian, Carol J Bult, Karl J Campbell, Riccardo Castiglia, Yung-Hao Ching, Amanda J Chunco, James J Crowley, Elissa J Chesler, John E French, Sofia I Gabriel, Daniel M Gatti, Theodore Garland, Eva B Giagia-Athanasopoulou, Mabel D Giménez, Sofia A Grize, Islam Gündüz, Andrew Holmes, Heidi C Hauffe, Jeremy S Herman, James M Holt, Kunjie Hua, Wesley J Jolley, Anna K Lindholm, María J López-Fuster, George Mitsainas, Maria da Luz Mathias, Leonard McMillan, M Graça Ramalhinho, Barbara Rehermann, Stephan P Rosshart, Jeremy B Searle, Meng-Shin Shiao, Emanuela Solano, Karen L Svenson, Pat Thomas-Laemont, David W Threadgill, Jacint Ventura, George M Weinstock, Daniel Pomp, Gary A Churchill, Fernando Pardo-Manuel de Villena

## Abstract

A selective sweep is the result of strong positive selection driving newly occurring or standing genetic variants to fixation, and can dramatically alter the pattern and distribution of allelic diversity in a population. Population-level sequencing data have enabled discoveries of selective sweeps associated with genes involved in recent adaptations in many species. In contrast, much debate but little empirical evidence addresses whether “selfish” genes are capable of fixation  thereby leaving signatures identical to classical selective sweeps  despite being neutral or deleterious to organismal fitness. We previously described *R2d2*, a large copy-number variant that causes non-random segregation of mouse Chromosome 2 in females due to meiotic drive. Here we show population-genetic data consistent with a selfish sweep driven by alleles of *R2d2* with high copy number (*R2d2^HC^*) in natural populations. We replicate this finding in multiple closed breeding populations from six outbred backgrounds segregating for *R2d2* alleles. We find that *R2d2^HC^* rapidly increases in frequency, and in most cases becomes fixed in significantly fewer generations than can be explained by genetic drift. *R2d2^HC^* is also associated with significantly reduced litter sizes in heterozygous mothers, making it a true selfish allele. Our data provide direct evidence of populations actively undergoing selfish sweeps, and demonstrate that meiotic drive can rapidly alter the genomic landscape in favor of mutations with neutral or even negative effects on overall Darwinian fitness. Further study will reveal the incidence of selfish sweeps, and will elucidate the relative contributions of selfish genes, adaptation and genetic drift to evolution.

## Introduction

Population-level sequencing data have enabled analyses of positive selection in many species, including mice (Staubach et al. 2012) and humans (Williamson et al. 2007; Grossman et al. 2013; Colonna et al. 2014). These studies seek to identify genetic elements, such as single nucleotide variants (SNVs) and copy number variants (CNVs), that are associated with phenotypic differences between populations that share a common origin (Fu and Akey 2013; Bryk and Tautz 2014). A marked difference in local genetic diversity between closely related taxa might indicate that one lineage has undergone a sweep, in which a variant under strong positive selection rises in frequency and carries with it linked genetic variation (“genetic hitch-hiking”), thereby reducing local haplotype diversity (Maynard Smith and Haigh 1974; Kaplan et al. 1989). In genomic scans for sweeps, it is typically assumed that the driving allele will have a strong positive effect on organismal fitness. Prominent examples of sweeps for which this assumption holds true (*i.e.* classic selective sweeps) include alleles at the *Vkorc1* locus, which confers rodenticide resistance in the brown rat (Pelz et al. 2005), and enhancer polymorphisms conferring lactase persistence in human beings (Bersaglieri et al. 2004). However, we and others have suggested that selfish alleles that strongly promote their own transmission irrespective of their effects on overall fitness could give rise to genomic signatures indistinguishable from those of classic selective sweeps (Sandler and Novitski 1957; White 1978; Henikoff and Malik 2002; Derome et al. 2004; Pardo-Manuel de Villena 2004; Brandvain and Coop 2011).

Suggestive evidence that sweeps may be driven by selfish alleles comes from studies in *Drosophila*. Incomplete sweeps have been identified at the *Segregation Distorter* (SD) locus (Presgraves et al. 2009) and in at least three X-chromosome systems (Babcock and Anderson 1996; Dyer et al. 2007; Derome et al. 2008; Kingan et al. 2010), all of which drive through the male germline. In addition, genomic conflict has been proposed as a possible driver of two nearly complete sweeps in *D. mauritiana* (Nolte et al. 2013). Incomplete sweeps were also detected in natural populations of *Mimulus* (monkeyflower), and the cause was identified as female meiotic drive of the centromeric *D* locus (Fishman and Saunders 2008). That all evidence of selfish sweeps derives from two genera is to some extent reflective of a selection bias, but may also indicate a difference in the incidence or effect of selfish alleles between these taxa and equally well-studied mammalian species (*e.g*. humans and mice). Furthermore, the lack of completed selfish sweeps reported in the literature may be due to an unexpected strength of balancing selection, in which the deleterious effects of selfish alleles prevent them from driving to fixation, or due to insufficient methods of detection.

Here, we investigate whether a selfish allele can sweep in natural and laboratory populations of the house mouse, *M. m. domesticus*. We recently described *R2d2*, a meiotic drive responder locus on mouse Chromosome 2 (Didion et al. 2015). *R2d2* is a variably sized copy number gain of a 127 kb core element that contains a single annotated gene, *Cwc22* (a spliceosomal protein). In females heterozygous for *R2d2*, transmission of an allele with high copy number (*R2d2^HC^*) relative to an allele with low copy number (*R2d2^LC^*) is a quantitative trait controlled by multiple unlinked modifier loci. We define “high copy number” as the minimum copy number with evidence of distorted transmission in existing experiments – approximately 7 units of the core element. Distorted transmission is present in some laboratory crosses (Siracusa et al. 1991; Montagutelli et al. 1996; Swallow et al. 1998; Eversley et al. 2010; Kelly et al. 2010a) segregating for *R2d2^HC^* alleles, but absent in others (Didion et al. 2015). *R2d2^HC^* genotype is also either uncorrelated or negatively correlated with litter size – a major component of absolute fitness – depending on the presence of meiotic drive. *R2d2^HC^* therefore behaves as a selfish genetic element. In the current study, we provide evidence of a recent sweep at *R2d2^HC^* in wild *M. m. domesticus* mice, and we show that *R2d2^HC^* has repeatedly driven selfish sweeps in closed-breeding mouse populations.

## Results and Discussion

### Evidence for a selfish sweep in wild mouse populations

A recent study (Pezer et al. 2015) showed extreme copy number variation at *Cwc22* in a sample of 26 wild mice (*M. m. domesticus*). To determine whether this was indicative of *R2d2* copy number variation in the wild, we assayed an additional 396 individuals sampled from 14 European countries and the United States (**Supplementary Table 1** and **Figure 1**). We found that *R2d2^HC^* alleles are segregating at a wide range of frequencies in natural populations (0.00 - 0.67; **Table 1**).

**Figure 1.**
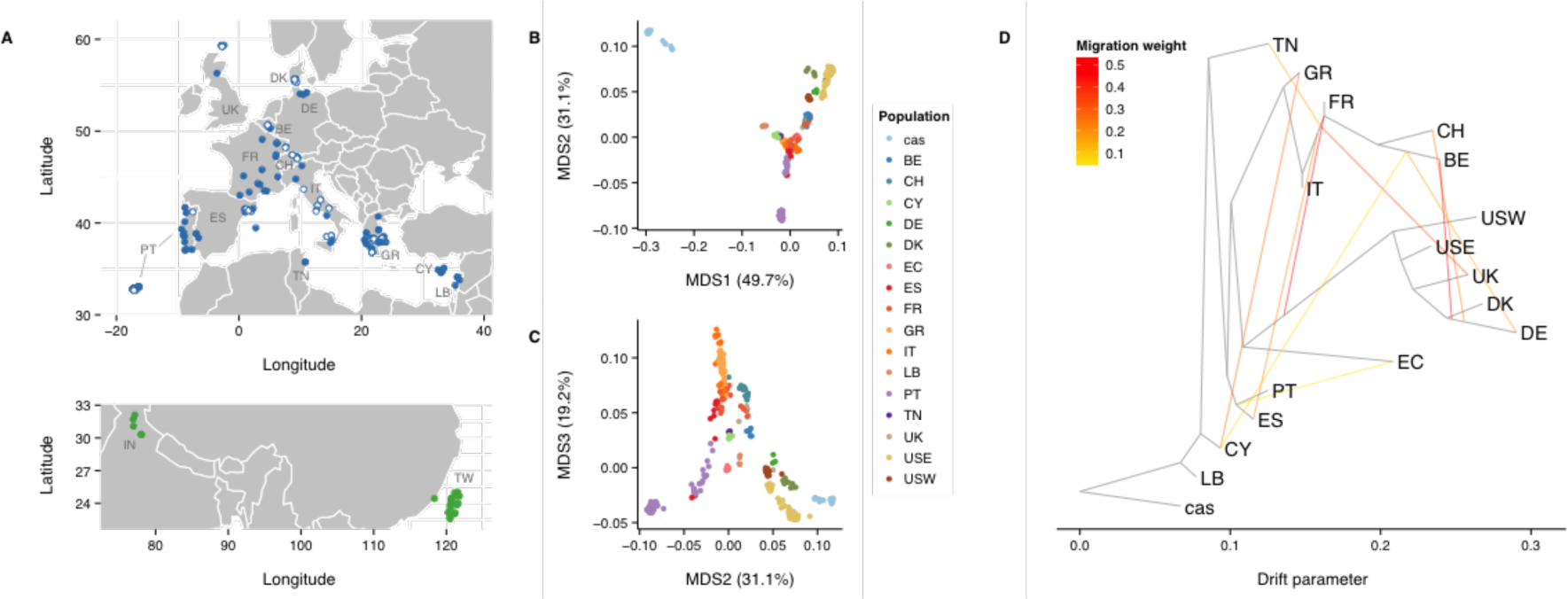
Wild mouse populations used in this study. **(A)** Geographic distribution of samples used in this study. Samples are colored by taxonomic origin: blue for *M. m. domesticus*, green for *M. m. castaneus*. Those with standard karyotype (2*n* = 40) are indicated by closed circles; samples with Robertsonian fusion karyotypes (2*n* < 40) are indicated by open circles. Populations from Floreana Island (Galapagos Islands, Ecuador; “EC”), Farallon Island (off the coast of San Francisco, California, United States; “USW"), and Maryland, United States (“USE”) are not shown. **(B,C)** Multidimensional scaling (MDS) (*k* = 3 dimensions) reveals population stratification consistent with geography. *M. m. domesticus* populations are labeled by country of origin. Outgroup samples of *M. m. castaneus* origin are combined into a single cluster (“cas”). **(D)** Population graph estimated from autosomal allele frequencies by TreeMix. Black edges indicate ancestry, while red edges indicate gene flow by migration or admixture. Topography of the population graph is consistent with MDS result and with the geographic origins of the samples.

**Table 1.** *R2d2^HC^* allele frequencies in wild *M. m. domesticus* populations.

| Population | R2d2HC Allele<br>Freq | 2 x (Number of<br>Individuals) |
| --- | --- | --- |
| BE | 0.50 | 6 |
| CH | 0.32 | 28 |
| CY | 0.00 | 14 |
| DE | 0.67 | 6 |
| DK | 0.06 | 18 |
| EC | 0.00 | 24 |
| ES | 0.22 | 18 |
| FR | 0.15 | 26 |
| GR | 0.08 | 106 |
| IT | 0.09 | 34 |
| LB | 0.25 | 8 |
| PT | 0.13 | 54 |
| TN | 0.00 | 4 |
| UK | 0.00 | 6 |
| USE | 0.21 | 102 |
| USW | 0.00 | 24 |

To test for a selfish sweep at *R2d2^HC^*, we genotyped the wild-caught mice on the MegaMUGA array (Rogala et al. 2014; Morgan and Welsh 2015; Morgan et al. 2015) and examined patterns of haplotype diversity. In the case of strong positive selection, unrelated individuals are more likely to share extended segments identical by descent (IBD) in the vicinity of the selected locus (Albrechtsen et al. 2010), compared with a population subject only to genetic drift. Consistent with this prediction, we observed an extreme excess of shared IBD across populations around *R2d2* (**Figure 2A**): *R2d2* falls in the top 0.25% of IBD-sharing scores across the autosomes. In all cases, the shared haplotype has high copy number, and this haplotype appears to have a single origin in European mice (**Supplementary Figure 1**). Strong signatures are also evident at a previously identified target of positive selection, the *Vkorc1* locus (distal Chromosome 7) (Song et al. 2011).

**Figure 2.**
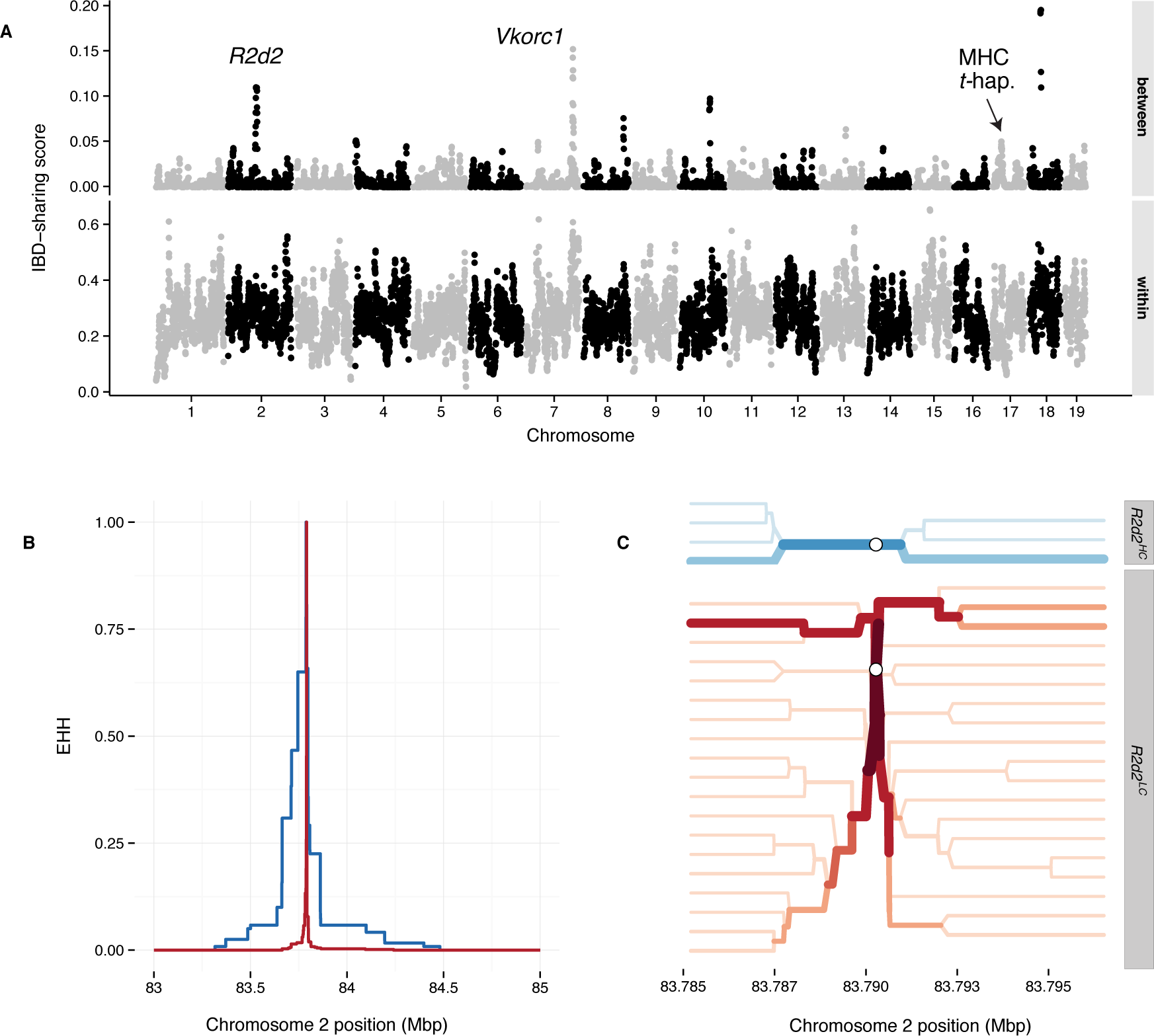
Haplotype-sharing at *R2d2* provides evidence of a selective sweep in wild mice of European origin. **(A)** Weighted haplotype-sharing score (see **Online Methods**), computed in 500 kb bins across autosomes, when those individuals are drawn from the same population (lower panel) or different populations (upper panel). Peaks of interest overlay *R2d2* (Chromosome 2; see **Supplementary Figure 2** for zoomed-in view) and *Vkorc1* (distal Chromosome 7). The position of the closely linked *t*-haplotype and MHC loci is also marked. **(B)** Decay of extended haplotype homozygosity (EHH) (Sabeti et al. 2002) on the *R2d2^HC^*-associated (blue) versus the *R2d2^LC^*-associated (red) haplotype. EHH is measured outward from the index SNP at chr2:83,790,275 and is bounded between 0 and 1. **(C)** Haplotype bifurcation diagrams for the *R2d2^HC^* (top panel, red) and *R2d2^LC^* (bottom panel, blue) haplotypes at the index SNP (open circle). Darker colors and thicker lines indicate higher haplotype frequencies. Haplotypes extend 100 sites in each direction from the index SNP.

In principle, the strength and age of a sweep can be estimated from the extent of loss of genetic diversity around the locus under selection. From the SNP data, we identified a ∼1 Mb haplotype with significantly greater identity between individuals with *R2d2^HC^* alleles compared to the surrounding sequence. We used published sequencing data from 26 wild mice (Pezer et al. 2015) to measure local haplotype diversity around *R2d2* and found that the haplotypes associated with *R2d2^HC^* alleles are longer than those associated with *R2d2^LC^* (**Figure 2B-C**). This pattern of extended haplotype homozygosity is consistent with positive selection over an evolutionary timescale as short as 450 generations (see **Methods**). However, due to the extremely low rate of recombination in the vicinity of *R2d2* (Didion et al. 2015), this is most likely an underestimate of the true age of the mutation.

It is important to note that the excess IBD we observe at *R2d2* (**Figure 2A**) arises from segments shared *between* geographically distinct populations (**Figure 1**). When considering sharing *within* populations only (**Supplementary Figure 2**), *R2d2* is no longer an outlier. Therefore, it was unsurprising that we failed to detect a sweep around *R2d2* using statistics that are designed to identify population-specific differences in selection, like hapFLK (Fariello et al. 2013), or selection in aggregate, like iHS (Voight et al. 2006) (**Supplementary Figure 3**).

**Figure 3.**
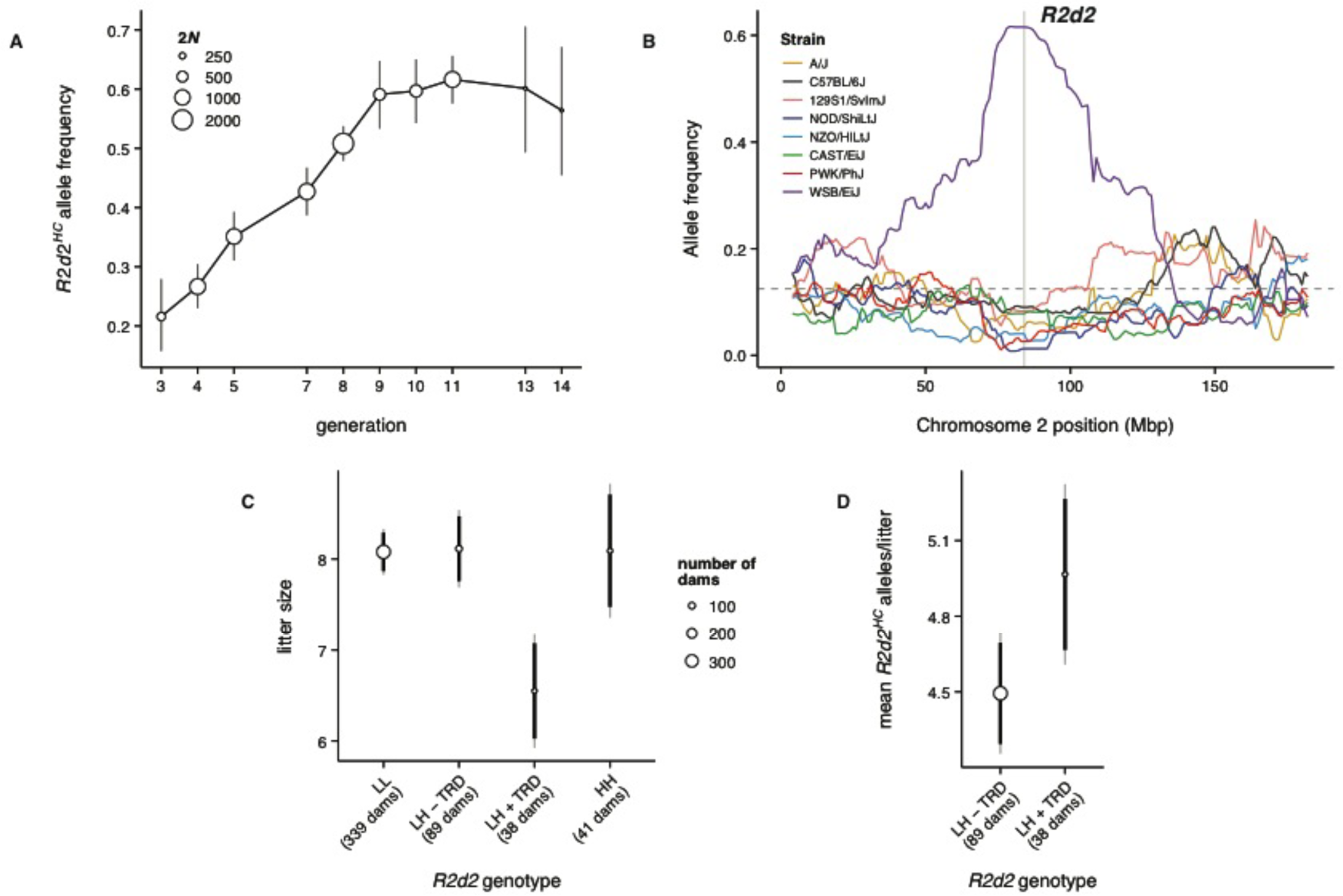
An *R2d2^HC^* allele rises to high frequency despite negative effect on litter size in the DO. **(A)** *R2d2* drives three-fold increase in WSB/EiJ allele frequency in 13 generations in the DO population. Circle sizes reflect number of chromosomes genotyped (2*N*); error bars are +/- 2 SE. **(B)** Allele frequencies across Chromosome 2 (averaged in 1 Mb bins) at generation 13 of the DO, classified by founder strain. Grey shaded region is the candidate interval for *R2d2*. **(C)** Mean litter size among DO females according to *R2d2* genotype: LL, *R2d2^LC/LC^*; LH – TRD, *R2d2^LC/HC^* without transmission ratio distortion; LH + TRD, *R2d2^LC/HC^* with transmission ratio distortion; HH, *R2d2^HC/HC^*. Circle sizes reflect number of females tested; error bars are 95% confidence intervals from a linear mixed model which accounts for parity and repeated measures on the same female (see **Methods**.) **(D)** Mean absolute number of *R2d2^HC^* alleles transmitted in each litter by heterozygous females with (LL + TRD) or without (LL – TRD) transmission ratio distortion. LL + TRD females transmit more *R2d2^HC^* alleles despite their significantly reduced litter size.

### A selfish sweep in an outbred laboratory population

We validated the ability of *R2d2^HC^* to drive a selfish sweep by examining *R2d2* allele frequencies in multiple closed-breeding laboratory populations for which we had access to samples from the founder populations. The Diversity Outbred (DO) is a randomized outbreeding population derived from eight inbred mouse strains that is maintained under conditions designed to minimize the effects of both selection and genetic drift (Svenson et al. 2012). Expected time to fixation or loss of an allele present in the founder generation (with initial frequency 1/8) is ∼900 generations. The WSB/EiJ founder strain contributed an *R2d2^HC^* allele which underwent a more than three-fold increase (from 0.18 to 0.62) in 13 generations (*p* < 0.001 by simulation; range 0.03 - 0.26 after 13 generations in 1000 simulation runs) (**Figure 3A**), accompanied by significantly distorted allele frequencies (p < 0.01 by simulation) across a ∼100 Mb region linked to the allele (**Figure 3B**).

### R2d2^HC^ has an underdominant effect on fitness

The fate of a selfish sweep depends on the fitness costs associated with the different genotypic classes at the selfish genetic element. For example, maintenance of intermediate frequencies of the *t*-complex (Lyon 1991) and *SD* (Hartl 1973) chromosomes in natural populations of mice and *Drosophila*, respectively, is thought to result from decreased fecundity associated with those selfish elements.

To assess the fitness consequences of *R2d2^HC^* we treated litter size as a proxy for absolute fitness (**Figure 3C**). We determined whether each female had distorted transmission of *R2d2* using a one-sided exact binomial test for deviation from the expected Mendelian genotype frequencies in her progeny. Average litter size among DO females homozygous for *R2d2^LC^* (“LL” in **Figure 3C**: 8.1; 95% CI 7.8 - 8.3) is not different than among females homozygous for *R2d2^HC^* (“HH”: 8.1; 95% CI 7.4 - 8.9) or among heterozygous females without distorted transmission of *R2d2^HC^* (“LH-TRD”: 8.1; 95% CI 7.7 - 8.5). However, in the presence of meiotic drive, litter size is markedly reduced (“LH+TRD”: 6.5; 95% CI 5.9 - 7.2; *p* = 5.7 × 10^−5^ for test of difference versus all other classes). The relative fitness of heterozygous females with distorted transmission is *w* = 0.81, for a selection coefficient of *s* = 1 – *w* = 0.19 (95% CI 0.10 – 0.23) against the heterozygote. Despite this underdominant effect, the absolute number of *R2d2^HC^* alleles transmitted by heterozygous females in each litter is significantly higher in the presence of meiotic drive than its absence (*p* = 0.032; **Figure 3D**). The rising frequency of *R2d2^HC^* in the DO thus represents a truly selfish sweep.

### Selfish sweeps in other laboratory populations

We also observed selfish sweeps in selection lines derived from the ICR:Hsd outbred population (Swallow et al. 1998), in which *R2d2^HC^* alleles are segregating (**Figure 4A**). Three of four lines selectively bred for high voluntary wheel-running (HR lines) and two of four control lines (10 breeding pairs per line per generation in both conditions) went from starting *R2d2^HC^* frequencies ∼0.75 to fixation in 60 generations or less: two lines were fixed by generation 20, and three more by generation 60. In simulations mimicking this breeding design and neutrality (**Figure 4B**), median time to fixation was 46 generations (5th percentile: 9 generations). Although the *R2d2^HC^* allele would be expected to eventually fix by drift in 6 of 8 lines given its high starting frequency, fixation in two lines within 20 generations and three more lines by 60 generations is not expected (*p* = 0.003 by simulation). In a related advanced intercross segregating for high and low copy number alleles at *R2d2* (HR8×C57BL/6J (Kelly et al. 2010b)), we observed that *R2d2^HC^* increased from a frequency of 0.5 to 0.85 in just 10 generations and fixed by 15 generations (**Figure 4C**), versus a median 184 generations in simulations (*p* < 0.001; **Figure 4D**). The increase in *R2d2^HC^* allele frequency in the DO and the advanced intercross populations occurred at least an order of magnitude faster than what is predicted by drift alone.

**Figure 4.**
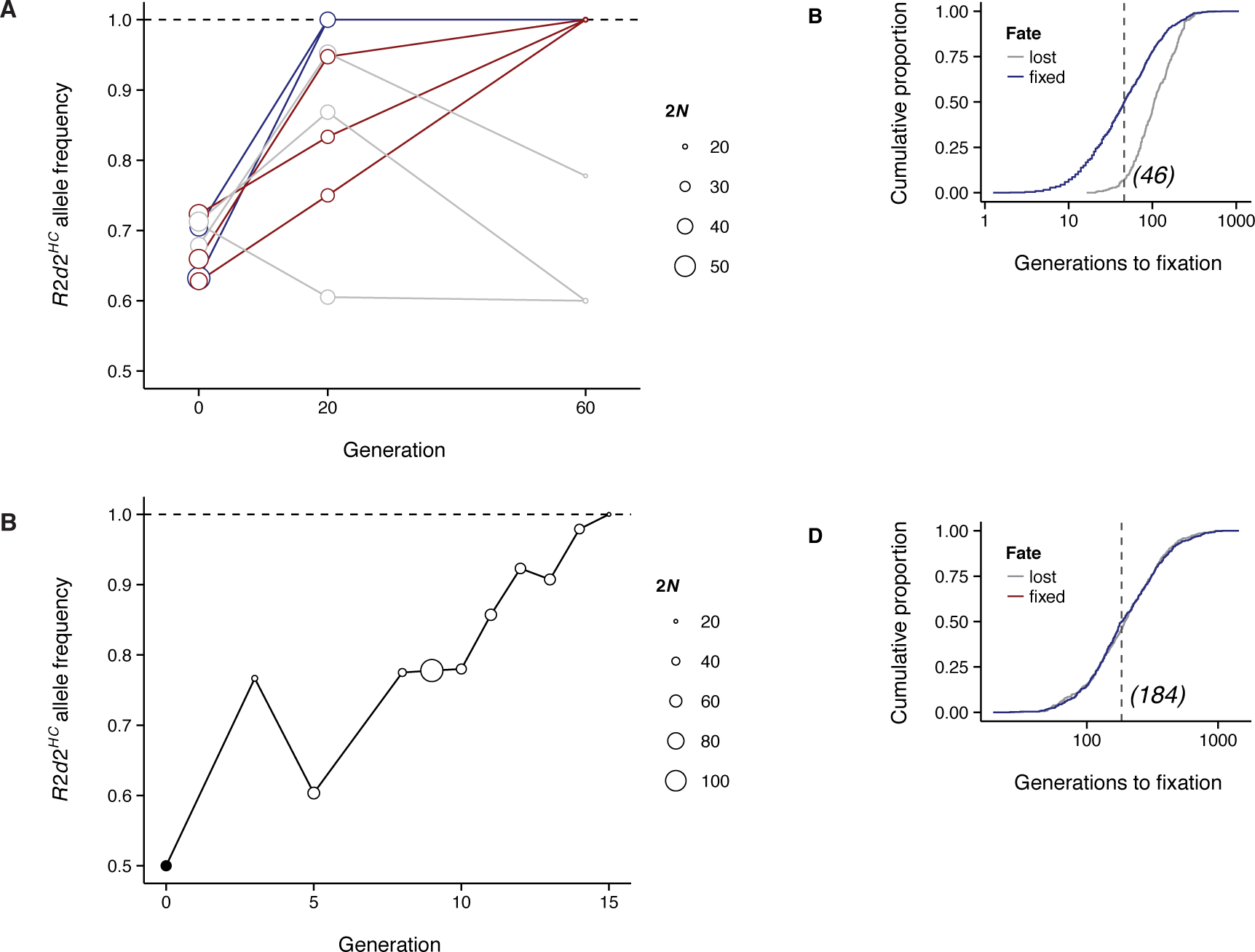
*R2d2^HC^* alleles rapidly increase in frequency in ICR:Hsd-derived laboratory populations. **(A)** *R2d2^HC^* allele frequency during breeding of 4 HR selection lines and 4 control lines. Trajectories are colored by their fate: blue, *R2d2^HC^* fixed by generation 20; red. *R2d2^HC^* fixed by generation 60; grey, *R2d2^HC^* not fixed. Circle sizes reflect number of chromosomes (*2N*) genotyped. **(B)** Cumulative distribution of time to fixation (blue) or loss (grey) of the focal allele in 1,000 simulations of an intercross line mimicking the HR breeding scheme. Dotted line indicates median fixation time. **(C)** *R2d2^HC^* allele frequency during breeding of an (HR8xC57BL/6J) advanced intercross line. Circle sizes reflect number of chromosomes (2*N*) genotyped. **(D)** Cumulative distribution of time to fixation (blue) or loss (grey) of the focal allele in 1,000 simulations of an advanced intercross line mimicking the HR8xC57BL/6J AIL. Dotted line indicates median fixation time.

Using archival tissue samples, we were able to determine *R2d2* allele frequencies in the original founder populations of 6 of the ∼60 wild-derived inbred strains available for laboratory use (Didion and Pardo-Manuel de Villena 2013). In four strains, WSB/EiJ, WSA/EiJ, ZALENDE/EiJ, and SPRET/EiJ, *R2d2^HC^* alleles were segregating in the founders and are now fixed in the inbred populations. In the other two strains, LEWES/EiJ and TIRANO/EiJ, the founders were not segregating for *R2d2* copy number and the inbred populations are fixed, as expected, for *R2d2^LC^* (**Supplementary Figure 4**). This trend in wild-derived strains is additional evidence of the tendency for *R2d2^HC^* to go to fixation in closed breeding populations when segregating in the founder individuals.

### On the distribution and frequency of R2d2^HC^ alleles in the wild

Considering the degree of transmission distortion in favor of *R2d2^HC^* (up to 95% (Didion et al. 2015)), and that *R2d2^HC^* repeatedly goes to fixation in laboratory populations, the moderate frequency of *R2d2^HC^* in the wild (0.14 worldwide, **Table 1**) is initially surprising. Neither do we find any obvious association between geography and *R2d2^HC^* allele frequency that might indicate the mutation’s origin or its pattern of gene flow (**Table 1** and **Figure 1**).

Several observations may explain these results. First, relative to the effective size of *M. m. domesticus* (82,500-165,000 (Geraldes et al. 2011)), our sample size was small. Our sampling was also geographically sparse and non-uniform. Thus, our allele frequency estimates may differ substantially from the true population allele frequencies at *R2d2*.

Second, the reduction in litter size associated with *R2d2^HC^* may have a greater impact on *R2d2* allele frequency in a natural population than in the controlled laboratory populations we studied. In these breeding schemes each mating pair contributes the same number of offspring to the next generation so that most fitness differences are effectively erased.

Third, *R2d2^HC^* alleles may be unstable and lose the ability to drive upon reverting to low copy number. This has been reported previously (Didion et al. 2015).

Fourth, in an large population (*i.e*. in the wild) the dynamics of an underdominant meiotic drive allele are only dependent on the relationship between the degree of transmission distortion (*m*) and the strength of selection against heterozygotes (Hedrick 1981) (*s*). This relationship can be expressed by the quantity *q* (see **Methods**), for which *q* > 1 indicates increasing probability of fixation of the driving allele, *q* < 1 indicates increasing probability that the allele will be purged, and *q* ≈ 1 leads to maintenance of the allele at an (unstable) equilibrium frequency. The fate of the driving allele in a finite population additionally depends on the population size (Hedrick 1981): the smaller the population, the greater the likelihood that genetic drift will fix a mutation with *q* < 1 (**Figure 5A-B**). We note that *R2d2^HC^* appears to exist close to the *q* ≈ 1 boundary (*s* ≈ 0.2, *m* ≈ 0.7, and thus *q* ≈ 0.96).

**Figure 5.**
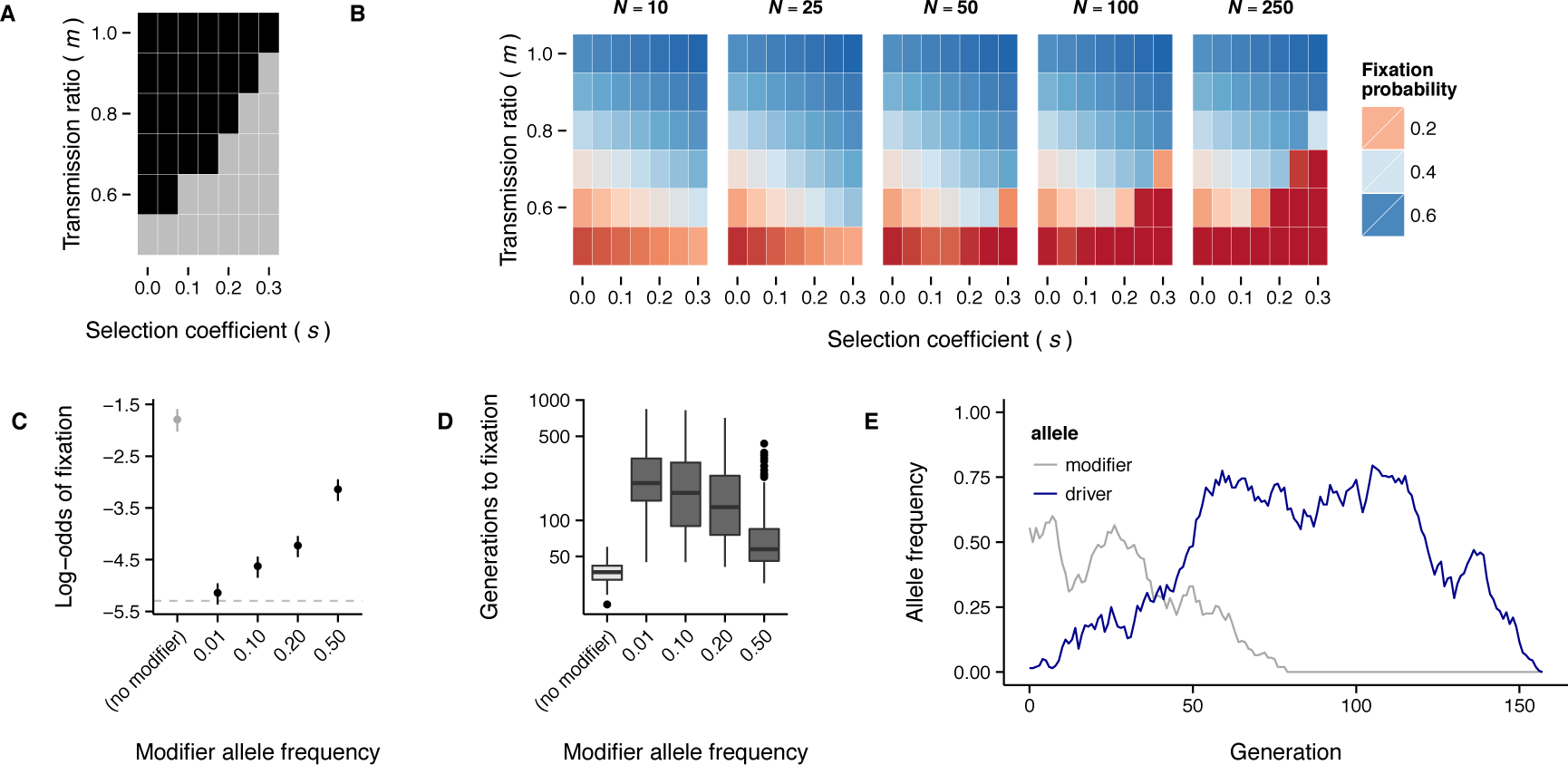
Population dynamics of a meiotic drive allele. **(A)** Phase diagram for a meiotic drive system like *R2d2*, with respect to transmission ratio (*m*) and selection coefficient against the heterozygote (*s*). Regions of the parameter space for which there is directional selection for the driving allele are shown in black; regions in which there are unstable equilibria or directional selection against the driving allele are shown in grey. **(B)** Probability of fixing the driving allele as a function of *m*, *s* and population size (*N*). Notice that, in the area corresponding to the grey region of panel A, fixation probability declines rapidly as population size increases. **(C)** Probability of fixing the driving allele in simulations of meiotic drive dependent on a single modifier locus (*N* = 100, *s* = 0.2, maximum *m* = 0.8, initial driver frequency 1/2*N*). Estimates are given +/- 2 SE. Grey dashed line corresponds to fixation probability for a neutral allele (1/2*N*). **(D)** Time to fixation of the driving allele. Values represent 100 fixation events in each condition. **(E)** Example allele-frequency trajectories from a “collapsed” selfish sweep: while the modifier allele is present at intermediate frequency, the driving allele sweeps to a frequency of ∼ 0.75. After the modifier allele is lost, the driver drifts out of the population as well.

Last but not least, in contrast to many meiotic drive systems, in which the component elements are tightly linked, the action of *R2d2^HC^* is dependent on unlinked modifier loci whose frequencies, modes of action, and effect sizes are unknown. It is therefore difficult to predict the effect of the modifiers on *R2d2* allele frequencies in the wild. We used forward-in-time simulations to explore the effect of a single unlinked modifier locus on fixation probability of a driving allele. Under an additive model (*m* = 0.80 for modifier genotype *AA*, 0.65 for genotype *Aa* and 0.50 for genotype *aa*), fixation probability is reduced and time to fixation is increased by the presence of the modifier locus (**Figure 5C-D**). As the modifier allele becomes more rare, fixation probability approaches the neutral expectation (1/2*N*, where *N* is population size). Importantly, the driving allele tends to sweep until the modifier allele is lost, and then drifts either to fixation or loss (**Figure 5E**). Drift at modifier loci thus creates a situation akin to selection in a varying environment – one outcome of which is balancing selection (Gillespie 2010). This is consistent with the maintenance of *R2d2^HC^* at intermediate frequencies in multiple populations separated by space and time, as we observe in wild mice.

### Concluding remarks

Most analyses of positive selection in the literature assume that the likelihood of a newly arising mutation becoming established, increasing in frequency and even going to fixation within a population is positively correlated with its effect on organismal fitness. Here, we have shown that a selfish genetic element has repeatedly driven sweeps in which the change in allele frequency and the effect on organismal fitness are decoupled. Our results suggest that evolutionary studies should employ independent evidence to determine whether loci implicated as drivers of selective sweeps are adaptive or selfish.

Although a selfish sweep has clear implications for such experimental populations as the DO and the Collaborative Cross (Didion et al. 2015), the larger evolutionary implications of selfish sweeps are less obvious. On the one hand, sweeps may be relatively rare, as appears to be the case for classic selective sweeps in recent human history (Hernandez et al. 2011). On the other hand, theory and comparative studies indicate that selfish genetic elements may be a potent force during speciation (White 1978; Hedrick 1981; Pardo-Manuel de Villena and Sapienza 2001; Henikoff and Malik 2002; Brandvain and Coop 2011). With the growing appreciation for the potential importance of non-Mendelian genetics in evolution, and the increasing tractability of population-scale genetic analyses, we anticipate that the effects of selfish elements such as *R2d2* in natural populations, including their contributions to events of positive selection, will soon be elucidated.

Improved understanding of the mechanism of meiotic drive at *R2d2* may also enable practical applications of selfish genetic elements. As demonstrated by the recent use of RNA-guided genome editing to develop gene drive systems in mosquitos and fruit flies (Esvelt et al. 2014; Gantz and Bier 2015; Hammond et al. 2015), experimental manipulation of chromosome segregation is now feasible. *R2d2* is an attractive option for the development of a mammalian gene drive system because, as we have shown here, it has already proven capable of driving to fixation in multiple independent genetic backgrounds. Furthermore, there are multiple unlinked modifiers of *R2d2* that, when identified, might be exploited for fine-grained manipulation of transmission ratios.

## Materials and Methods

### Mice

Wild *M. m. domesticus* mice were trapped at a large number of sites across Europe and the Americas (**Figure 1A** (upper panel), **and Supplementary Table 1**). A set of 29 *M. m. castaneus* mice trapped in northern India and Taiwan (**Figure 1A**, lower panel) were included as an outgroup (Yang et al. 2011). Trapping was carried out in concordance with local laws, and either did not require approval or was carried out with the approval of the relevant regulatory bodies (depending on the locality and institution).

All Diversity Outbred (DO) mice were bred at The Jackson Laboratory. Litter sizes were counted within 24 hours of birth. Individual investigators purchased mice for unrelated studies and contributed either tissue samples or genotype data to this study (**Supplementary Table 2**).

High running (HR) selection and intercross lines were developed as previously described (Swallow et al. 1998; Kelly et al. 2010a; Leamy et al. 2012). Mouse tails were archived from 3 generations of the HR selection lines (-2, +22, and +61) and from every generation of the HR8xC57BL/6J advanced intercross.

Progenitors of wild-derived strains have various origins (see **Supplementary Methods**), and were sent to Eva M. Eicher at The Jackson Laboratory for inbreeding in the early 1980s. Frozen tissues from animals in the founder populations were maintained at The Jackson Laboratory by Muriel Davidson until 2014, when they were transferred to the Pardo-Manuel de Villena laboratory at the University of North Carolina at Chapel Hill.

All laboratory mice were handled in accordance with the IACUC protocols of the investigators’ respective institutions.

### Genotyping

*Microarray genotyping and quality control:* Whole-genomic DNA was isolated from tail, liver, muscle or spleen using Qiagen Gentra Puregene or DNeasy Blood & Tissue kits according to the manufacturer’s instructions. All genome-wide genotyping was performed using the Mouse Universal Genotyping Array (MUGA) and its successor, MegaMUGA (GeneSeek, Lincoln, NE) (Collaborative Cross Consortium 2012; Morgan and Welsh 2015). Genotypes were called using Illumina BeadStudio (Illumina Inc., Carlsbad, CA). We excluded all markers and all samples with missingness greater than 10%. We also computed the sum intensity for each marker: 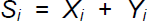, where *X_i_* and *Y_i_* are the normalized hybridization intensities of the two allelic probes. We determined the expected distribution of sum intensity values using a large panel of control samples. We excluded any array for which the set of intensities 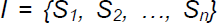 was not normally distributed or whose mean was significantly left-shifted from the reference distribution (one-tailed *t*-test with *p* < 0.05).

*PCR genotyping:* The *R2d2* element has been mapped to a 900 kb critical region on Chromosome 2: 83,631,096 - 84,541,308 (mm9 build), referred to herein as the “candidate interval” (Didion et al. 2015). We designed primers to amplify two regions within the candidate interval. *Primer Set A* targets a 318 bp region (chr2: 83,673,604 - 83,673,921) with two distinct haplotypes in linkage with either the *R2d2^LC^* allele or the *R2d2^HC^* allele: 5’-CCAGCAGTGATGAGTTGCCATCTTG-3’ (forward) and 5’- TGTCACCAAGGTTTTCTTCCAAAGGGAA-3’ (reverse). *Primer Set B* amplifies a 518 bp region (chr2: 83,724,728 - 83,725,233); the amplicon is predicted, based on whole-genome sequencing, to contain a 169 bp deletion in HR8 relative to the C57BL/6J reference genome: 5’-GAGATTTGGATTTGCCATCAA-3’ (forward) and 5’-GGTCTACAAGGACTAGAAACAG-3’ (reverse). Primers were designed using IDT PrimerQuest (https://www.idtdna.com/Primerquest/Home/Index).

Crude whole-genomic DNA for PCR reactions was extracted from mouse tails. The tissues were heated in 100 µl of 25 mM NaOH/0.2 mM EDTA at 95°C for 60 minutes followed by the addition of 100 µl of 40 mM Tris-HCl. The mixture was then centrifuged at 2000 × *g* for 10 minutes and the supernatant used as PCR template. PCR reactions contained 1 µL dNTPs, 0.3 µL of each primer, 5.3 µL of water, and 0.1 µL of GoTaq polymerase (Promega) in a final volume of 10 µL. Cycling conditions were 95°C, 2-5 min, 35 cycles at 95°, 55° and 72°C for 30 sec each, with a final extension at 72°C, 7 min.

For *Primer Set A*, products were sequenced at the University of North Carolina Genome Analysis Facility on an Applied Biosystems 3730XL Genetic Analyzer. Chromatograms were analyzed with the Sequencher software package (Gene Codes Corporation, Ann Arbor, Michigan, United States). For *Primer Set B*, products were visualized and scored on 2% agarose gels. Assignment to haplotypes was validated by comparing the results to qPCR assays for the single protein-coding gene within *R2d2*, *Cwc22* (see “Copy-number assays” below). For generation +61, haplotypes were assigned based on MegaMUGA genotypes and validated by the normalized per-base read depth from whole-genome sequencing (see below), calculated with samtools mpileup (Li et al. 2009). The concordance between qPCR, read depth, and haplotypes assigned by MegaMUGA or Sanger sequencing is shown in **Supplementary Figure 5.**

*Assays:* Wild mice were genotyped on MegaMUGA (**Supplementary Table 1**). DO mice were genotyped on MUGA and MegaMUGA (**Supplementary Table 2**). HR selection lines were genotyped at three generations, one before (-2) and two during (+22 and +61) artificial selection. We genotyped 185 randomly selected individuals from generation -2 and 157 individuals from generation +22 using *Primer Set A*. An additional 80 individuals from generation +61 were genotyped with the MegaMUGA array (see “Microarray genotyping and quality-control” below). The HR8xC57BL/6J advanced intercross line was genotyped with *Primer Set B* in tissues from breeding stock at generations 3, 5, 8, 9, 10, 11, 12, 13, 14, and 15.

***Copy-number assays and assignment of* R2d2 *status***. Copy-number at *R2d2* was determined by qPCR for *Cwc22*, the single protein-coding gene in the *R2d* repeat unit, as described in detail in (Didion et al. 2015). Briefly, we used commercially available TaqMan kits (Life Technologies assay numbers Mm00644079_cn and Mm00053048_cn) to measure the copy number of *Cwc22* relative to the reference genes *Tfrc* (cat. no. 4458366, for target Mm00053048_cn) or *Tert* (cat. no. 4458368, for target Mm00644079_cn). Cycle thresholds (*C*_t_) were determined for each target using ABI CopyCaller v2.0 software with default settings, and relative cycle threshold was calculated as

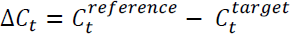

We normalized the ∆*C_t_* across batches by fitting a linear mixed model with batch and target-reference pair as random effects.

Estimation of integer diploid copy numbers > ~3 by qPCR is infeasible without many technical and biological replicates, especially in the heterozygous state. We took advantage of *R2d2* diploid copy-number estimates from whole-genome sequencing for the inbred strains C57BL/6J (0), CAST/EiJ (2) and WSB/EiJ (66), and the (WSB/EiJxC57BL/6J)F_1_ (33) to establish a threshold for declaring a sample “high-copy.” For each of the two TaqMan target-reference pairs we calculated the sample mean (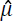) and standard deviation (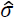) of the normalized ∆*C_t_* among CAST/EiJ controls and wild *M. m. castaneus* individuals together. We designated as “high-copy” any individual with normalized ∆*C_t_* greater than 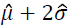 - that is, any individual with approximately > 95% probability of having diploid copy number >2 at *R2d2*. Individuals with high copy number and evidence of local heterozygosity (a heterozygous call at any of the 13 markers in the *R2d2* candidate interval) were declared heterozygous *R2d2^HC/LC^*, and those with high copy number and no heterozygous calls in the candidate interval were declared homozygous *R2d2^HC/HC^*.

### Exploration of population structure in wild mice

Scans for signatures of positive selection based on patterns of haplotype-sharing assume that individuals are unrelated. We identified pairs of related individuals using the *IBS2\** ratio (Stevens et al. 2011), defined as *HETHET* / (*HOMHOM + HETHET)*, where *HETHET* and *HOMHOM* are the count of non-missing markers for which both individuals are heterozygous (share two alleles) and homozygous for opposite alleles (share zero alleles), respectively. Pairs with *IBS2*\* < 0.75 were considered unrelated. Among individuals which were a member of one or more unrelated pairs, we iteratively removed one sample at a time until no related pairs remained, and additionally excluded markers with minor-allele frequency < 0.05 or missingness > 0.10. The resulting dataset contains genotypes for 396 mice at 58,283 markers.

Several of our analyses required that samples be assigned to populations. Because mice in the wild breed in localized demes and disperse only over short distances (on the order of hundreds of meters) (Pocock et al. 2005), it is reasonable to delineate populations on the basis of geography. We assigned samples to populations based on the country in which they were trapped. To confirm that these population labels correspond to natural clusters we performed two exploratory analyses of population structure. First, classical multidimensional scaling (MDS) of autosomal genotypes was performed with PLINK (Purcell et al. 2007) (—mdsplot—autosome). The result is presented in **Figure 1B-C**, in which samples are colored by population. Second, we used TreeMix (Pickrell and Pritchard 2012) to generate a population tree allowing for gene flow using the set of unrelated individuals. Autosomal markers were first pruned to reach a set in approximate linkage equilibrium (plink—indep 25 1). TreeMix was run on the resulting set using the *M. m. castaneus* samples as an outgroup and allowing up to 10 gene-flow edges (treemix-root “cas” -k 10) (**Figure 1D**). The clustering of samples by population evident by MDS and the absence of long-branch attraction in the population tree together indicate that our choices of population labels are biologically reasonable.

***Scans for selection in wild mice***. Two complementary statistics, hapFLK (Fariello et al. 2013) and standardized iHS score (Voight et al. 2006), were used to examine wild-mouse genotypes for signatures of selection surrounding *R2d2*. The hapFLK statistic is a test of differentiation of local haplotype frequencies between hierarchically-structured populations. It can be interpreted as a gene ralization of Wright’s *F*_ST_ which exploits local LD. Its model for haplotypes is that of fastPHASE (Scheet 2006) and requires a user-specified value for the parameter *K*, the number of local haplotype clusters. We computed hapFLK in the set of unrelated individuals using *M. m. castaneus* samples as an outgroup for *K* = {4, 8, 12, 16, 20, 24, 28, 32} (hapflk—outgroup “cas” -k {K}) and default settings otherwise.

The iHS score (and its allele-frequency-standardized form |iHS|) is a measure of extended haplotype homozygosity on a derived haplotype relative to an ancestral one. For consistency with the hapFLK analysis, we used fastPHASE on the same genotypes over the same range of *K* with 10 random starts and 25 iterations of expectation-maximization (fastphase -K{K} -T10 -C25) to generate phased haplotypes. We then used selscan (Szpiech and Hernandez 2014) to compute iHS scores (selscan—ihs) and standardized the scores in 25 equally-sized bins (selscan-norm—bins 25).

Values in the upper tail of the genome-wide distribution of hapFLK or |iHS| represent candidates for regions under selection. We used percentile ranks directly and did not attempt to calculate approximate or empirical *p*-values.

***Detection of identity-by-descent (IBD) in wild mice***. As an alternative test for selection, we computed density of IBD-sharing using the RefinedIBD algorithm of BEAGLE v4.0 (r1399) (Browning and Browning 2013), applying it to the full set of 500 individuals. The haplotype model implemented in BEAGLE uses a tuning parameter (the “scale” parameter) to control model complexity: larger values enforce a more parsimonious model, increasing sensitivity and decreasing computational cost at the expense of accuracy. The authors recommend a value of 2.0 for ∼1M SNP arrays in humans. We increased the scale parameter to 5.0 to increase detection power given (a) our much sparser marker set, and (b) the relatively weaker local LD in mouse versus human populations (Laurie et al. 2007). We trimmed one marker from the ends of candidate IBD segments to reduce edge effects (java -jar beagle.jar ibd=true ibdscale=5 ibdtrim=1). We retained those IBD segments shared between individuals in the set of 396 unrelated mice. In order to limit noise from false-positive IBD segments, we further removed segments with LOD score < 5.0 or width < 0.5 cM.

An empirical IBD-sharing score was computed in 500 kb bins with 250 kb overlap as:

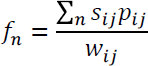

where the sum in the numerator is taken over all IBD segments overlapping bin *n* and *s_ij_* is an indicator variable which takes the value 1 if individuals *i,j* share a haplotype IBD in bin *n* and 0 otherwise. The weighting factor *w_ij_* is defined as

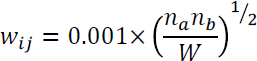

with

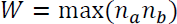

where *n_a_* and *n_b_* are the number of unrelated individuals in the population to which individuals *i* and *j* belong, respectively. This weighting scheme accounts for the fact that we oversample some geographic regions (for instance, Portugal and Maryland) relative to others. To explore differences in haplotype-sharing within versus between populations we introduce an additional indicator *p_ij_*. Within-population sharing is computed by setting *p_ij_* = 1 if individuals *i,j* are drawn from the same population and *p_ij_* = 0 otherwise. Between-population sharing is computed by reversing the values of *p_ij_*. The result is displayed in **Figure 2**.

***Analysis of local sequence diversity in whole-genome sequence from wild mice***. We obtained raw sequence reads for 26 unrelated wild mice (European Nucleotide Archive project accession PRJEB9450 (Pezer et al. 2015)); samples are listed in **Supplementary Table 3**. Details of the sequencing protocol are given in the indicated reference. Briefly, paired-end libraries with mean insert size 230 bp were prepared from genomic DNA using the Illumina TruSeq kit. Libraries were sequenced on the Illumina HiSeq 2000 platform with 2×100bp reads to an average coverage of 20X per sample (populations AHZ, CLG and FRA) or 12X per sample (population HGL). We realigned the raw reads to the mouse reference genome (GRCm38/mm10 build) using BWA MEM (Li and Durban, unpublished) with default parameters. SNPs relative to the reference sequence of Chromosome 2 were called using samtools mpileup v0.1.19-44428cd with maximum per-sample depth of 200. Genotype calls with root-mean-square mapping quality < 30 or genotype quality > 20 were treated as missing. Sites were used for phasing if they had a minor-allele count ≥ 2 and at most 2 missing calls. BEAGLE v4.0 (r1399) was used to phase the samples conditional on each other, using 20 iterations for phasing and default settings otherwise (java -jar beagle.jar phasing-its=20). Sites were assigned a genetic position by linear interpolation on the most recent genetic map for the mouse (Liu et al. 2010; Liu et al. 2014).

The *R2d2* candidate interval spans positions 83,790,939 - 84,701,151 in the mm10 reference sequence. As the index SNP for *R2d2^HC^* we chose the SNP with strongest nominal association with *R2d2* copy number (as estimated by Pezer *et al*. (2015)) within 1 kb of the proximal boundary of the candidate interval. That SNP is chr2:83,790,275T>C. The C allele is associated with high copy number and is therefore presumed to be the derived allele. We computed the extended haplotype homozygosity (EHH) statistic (Sabeti et al. 2002) in the phased dataset over a 1 Mb window on each side of the index SNP using selscan (selscan—ehh—ehh-win 1000000). The result is presented in **Figure 2B**. Decay of haplotypes away from the index SNP was visualized as a bifurcation diagram (**Figure 2C**) using code adapted from the R package rehh (https://cran.r-project.org/package=rehh).

***Estimation of age of* R2d2^HC^ *alleles in wild mice***. To obtain a lower bound for the age of *R2d2^HC^* and its associated haplotype, we used the method of Stephens *et al*. (1998) (Stephens et al. 1998). Briefly, this method approximates the probability *P* that a haplotype is affected by recombination or mutation during the *G* generations since its origin as

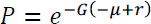

where *μ* and *r* are the per-generation rates of mutation and recombination, respectively. Assuming *μ* << *r* and, taking *P’*, the observed number of ancestral (non-recombined) haplotypes in a sample, as an estimator of *P*, obtain the following expression for *G*:

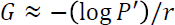

We enumerated haplotypes in our sample of 52 chromosomes at 3 SNPs spanning the *R2d2* candidate interval. The most proximal SNP is the index SNP for the EHH analyses (chr2:83,790,275T>C); the most distal SNP is the SNP most associated with copy number within 1 kbp of the boundary of the candidate interval (chr2:84,668,280T>C); and the middle SNP was randomly-chosen to fall approximately halfway between (chr2:84,079,970C>T). The three SNPs span genetic distance 0.154 cM (corresponding to *r* = 0.00154). The most common haplotype among samples with high copy number according to Pezer et al. (2015) was assumed to be ancestral. Among 52 chromosomes, 22 carried at least part of the *R2d2^HC^*-associated haplotype; of those, 11 were ancestral and 11 recombinant (**Supplementary Table 3**). This gives an estimated age of 450 generations for *R2d2^HC^*.

We note that the approximations underlying this model assume constant population size and neutrality. To the extent that haplotype homozygosity decays more slowly on a positively- (or selfishly-) selected haplotype, we will underestimate the true age of *R2d2^HC^*.

***Inference of local phylogeny at R2d2***. To determine whether the *R2d2^HC^* haplotype(s) shared among wild mice have a single origin, we constructed a phylogenetic tree from the 39 MegaMUGA SNPs in the region flanking *R2d2* (Chromosome 2: 82 - 85 Mb). We first excluded individuals heterozygous in the region and then constructed a matrix of pairwise distances from the proportion of alleles shared identical-by-state (IBS) between samples. A tree was inferred from the distance matrix using the neighbor-joining method implemented in the R package ape (http://cran.r-project.org/package=ape).

***Haplotype frequency estimation in the Diversity Outbred***. We inferred the haplotypes of DO individuals using probabilistic methods (Liu et al. 2010; Liu et al. 2014). We combined the haplotypes of DO individuals genotyped in this study with the Generation 8 individuals in Didion *et al*. (2015). As an additional QC step, we computed the number of historical recombination breakpoints per individual per generation (Svenson et al. 2012) and removed outliers (more than 1.5 standard deviations from the mean). We also excluded related individuals based on the distribution of haplotype sharing between related and unrelated individuals computed from simulations (mean 0.588 ± 0.045 for first-degree relatives; mean 0.395 ± 0.039 for second-degree relatives; and mean 0.229 ± 0.022 for unrelated individuals; see **Supplementary Methods**). Finally, we computed in each generation the frequency of each founder haplotype at 250 kb intervals surrounding the *R2d2* region (Chromosome 2: 78-86 Mb), and identified the greatest WSB/EiJ haplotype frequency.

***Analyses of fitness effects of* R2d2^HC^ *in the Diversity Outbred***. To assess the consequences of *R2d2^HC^* for organismal fitness, we treated litter size as a proxy for absolute fitness. Using breeding data from 475 females from DO generations 13, 16, 18 and 19, we estimated mean litter size in four genotype groups: females homozygous *R2d2^LC/LC^*; females heterozygous *R2d2^HC/LC^* with transmission ratio distortion (TRD) in favor of the *R2d2^HC^* allele; females heterozygous *R2d2^HC/LC^* without TRD; and females homozygous *R2d2^HC/HC^*. The 126 heterozygous females were originally reported in Didion et al. (2015). Group means were estimated using a linear mixed model with parity and genotype as fixed effects and a random effect for each female using the lme4 package for R. Confidence intervals were obtained by likelihood profiling and post-hoc comparisons were performed via *F*-tests, using the Kenward-Roger approximation for the effective degrees of freedom. The mean number of *R2d2^HC^* alleles transmitted per litter by heterozygous females with and without TRD was estimated from data in Didion et al. (2015) with a weighted linear model, using the total number of offspring per female as weights. Litter sizes are presented in **Supplementary Table 2**, and estimates of group mean litter sizes in **Figure 3C**.

***Whole-genome sequencing of HR selection lines***. Ten individuals from generation +61 of each of the eight HR selection lines were subject to whole-genome sequencing. Briefly, high-molecular-weight genomic DNA was extracted using a standard phenol/chloroform procedure. Illumina TruSeq libraries were constructed using 0.5 μg starting material, with fragment sizes between 300 and 500 bp. Each library was sequenced on one lane of an Illumina HiSeq2000 flow cell in a single 2×100bp paired-end run.

***Null simulations of closed breeding populations***. Widespread fixation of alleles due to drift is expected in small, closed populations such as the HR lines or the HR8·C57BL/6J advanced intercross line. But even in these scenarios, an allele under positive selection is expected to fix 1) more often than expected by drift alone in repeated breeding experiments using the same genetic backgrounds, and 2) more rapidly than expected by drift alone. We used the R package simcross (https://github.com/kbroman/simcross) to obtain the null distribution of fixation times and fixation probabilities for an HR line under Mendelian transmission.

We assume that the artificial selection applied for voluntary exercise in the HR lines (described in Swallow *et al*. (1998)) was independent of *R2d2* genotype. This assumption is justified for two reasons. First, 3 of 4 selection lines and 2 of 4 control (unselected) lines fixed *R2d2^HC^*. Second, at the fourth and tenth generation of the HR8xC57BL/6J advanced intercross, no quantitative trait loci (QTL) associated with the selection criteria (total distance run on days 5 and 6 of a 6-day trial) were found on Chromosome 2. QTL for peak and average running speed were identified at positions linked to *R2d2*; however, HR8 alleles at those QTL were associated with decreased, not increased, running speed (Kelly et al. 2010a; Leamy et al. 2012).

Without artificial selection an HR line reduces to an advanced intercross line maintained by avoidance of sibling mating. We therefore simulated 100 replicates of an advanced intercross with 10 breeding pairs and initial focal allele frequency 0.75. Trajectories were followed until the focal allele was fixed or lost. As a validation we confirmed that the focal allele was fixed in 754 of 1000 runs, not different from the expected 750 (*p* = 0.62, binomial test). Simulated trajectories and the distribution of sojourn times are presented in **Figure 4B**.

The HR8xC57BL/6J advanced intercross line was simulated as a standard biparental AIL with initial focal allele frequency of 0.5. Again, 1000 replicates of an AIL with 20 breeding pairs were simulated and trajectories were followed until the focal allele was fixed or lost. The result is presented in **Figure 4D**.

***Investigation of population dynamics of meiotic drive***. We used two approaches to investigate the population dynamics of a female-limited meiotic drive system with selection against the heterozygote. First, we evaluated the fixation probability of a driving allele in relationship to transmission ratio (*m*), selection coefficient against the heterozygote (*s*) and population size (*N*) by modeling the population as a discrete-time Markov chain whose states are possible counts of the driving allele. Define *p*_*t* + 1_ to be the expected frequency of the driving allele in generation *t*+1 given its frequency in the previous generation (*p_t_*). Following Hedrick et al. (1981), the expression for *p*_*t* + 1_ is

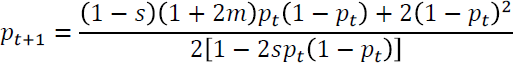

In an infinite population, the equilibrium behavior of the system is governed by the quantity *q*:

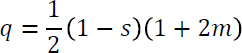

When *q* > 1, the driving allele always increases in frequency. For values of *q* ≈ 1 and smaller, the driving allele is either lost or reaches an unstable equilibrium frequency determined *m* and *s*.

Let *M* be the matrix of transition probabilities for the Markov chain with 2*N* + 1 states corresponding to possible counts of the driving allele in the population (0, … 2*N*). The entries *m_ij_* of *M* are

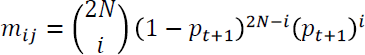

Given a vector *p*_0_ of starting probabilities, the probability distribution at generation *t* is obtained by iteration:

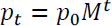

We initiated the chain with a single copy of the driving allele (i.e. *p*_0_ [1] = 1). Since this Markov chain has absorbing states (namely allele counts 0 and 2*N*), we approximated steady-state probabilities by iterating the chain until the change in probabilities between successive generations was < 10^−4^. Fixation probability is given by the value of the entry *p*_*t*_[2*N*] at convergence. We evaluated all possible combinations of 0.5 ≤ *m* ≤ 1.0 (in steps of 0.1) and 0 ≤ *s* ≤ 0.3 (in steps of 0.05).

To investigate the effects of modifier loci on the frequency trajectory of a driving allele we used forward-in-time simulations under a Wright-Fisher model with selection, implemented in Python. Simulations assumed a constant population size of 2*N* = 200 chromosomes, each 100 cM long, with balanced sex ratio. At the beginning of each run a driving allele was introduced (at 50 cM) on a single, randomly chosen chromosome. Modifier alleles were introduced into the population independently at a specified frequency, at position 0.5 cM (ie. unlinked to the driving allele). To draw the next generation, an equal number of male and female parents were selected (with replacement) from the previous generation according to their fitness. Among females heterozygous for the driving allele, transmission ratio (*m*) was calculated according to genotype at the modifier loci (if any). For males and homozygous females, *m* = 0.5. Individuals were assigned a relative fitness of 1 if *m* = 0.5 and 0.8 if *m* > 0.5. Recombination was simulated under the Haldane model (*i.e*. a Poisson process along chromosomes with no crossover interference). Finally, for each individual in the next generation, one chromosome was randomly chosen from each parent with probability *m*.

Simulation runs were restarted when the driving allele was fixed or lost, until 100 fixation events were observed in each condition of interest. Probability of fixation was estimated using the waiting time before each fixation event, assuming a geometric distribution of waiting times, using the fitdistr() function in the R package MASS. Simulations are summarized in **Figure 5**.

Simulation code is available on GitHub: https://github.com/andrewparkermorgan/r2d2-selfish-sweep.

## Acknowledgements

We wish to thank all the scientists and research personnel who collected and processed the samples used in this study. In particular we acknowledge Luanne Peters and Alex Hong-Tsen Yu for providing critical samples; Ryan Buus and T. Justin Gooch for isolating DNA for high-density genotyping of wild-caught mice; and Vicki Cappa, A. Cerveira, Daniel Förster, Guila Ganem, Ron and Annabelle Lesher, K. Saïd, Toni Schelts, Dan Small, and J. Tapisso for aiding in mouse trapping. We thank Muriel Davisson at the Jackson Laboratory for maintaining, for several decades, tissue samples from breeding colonies used to generate wild-derived inbred strains. We also thank Francis Collins, Jim Evans, Matthew Hahn, and Corbin Jones for comments on an earlier version of this manuscript. This work was supported by the National Institutes of Health T32GM067553 to JPD and APM, F30MH103925 to APM, P50GM076468 to EJC, GAC, and FPMV, K01MH094406 to JJC, DK-076050 and DK-056350 to DP, AG038070 to GAC, and the intramural research program to BR and SPR; National Science Foundation IOS-1121273 to TG; Vaadia-BARD Postdoctoral Fellowship Award FI-478-13 to LY; U.S. Army Medical Research and Materiel Command W81XWH-11-1-0762 to CJB; The Jackson Laboratory new investigator funds to EJC; The National Center for Scientific Research, France to JBD (This is contribution n°ISEM 2016-002); the University of Rome “La Sapienza” to RC and ES; Claraz-Stiftung to AL; Natural Environment Research Council (UK) to MDG, HCH and JBS; EU Human Capital and Mobility Programme (CHRX-CT93-0192) to HCH and JBS; Foundation for Science and Technology, Portugal PTDC/BIA-EVF/116884/2010 and UID/AMB/50017/2013 to SIG, MLM and JBS; Spanish “Ministerio de Ciencia y Tecnología” CGL2007-62111 and “Ministerio de Economía y Competitividad” CGL2010-15243 to JV; and the Oliver Smithies Investigator funds provided by the School of Medicine at University of North Carolina to FPMV.

## Author Contributions

JPD, GAC and FPMV conceived the study. JBD, CJB, KJC, RC, Y-HC, AJC, JJC, EJC, JEF, SIG, DMG, TG, EBG-A, MDG, SAG, IG, AH, HCH, JSH, JMH, KH, WJJ, AKL, MJL-F, GM, MM, LM, MGR, BR, SPR, JBS, MSS, ES, KLS, PT-L, DWT, JVQ, GMW, DP, GAC, and FPMV provided biological samples and/or unpublished data sets. APM, LY, TAB, RCM, and LOdS conducted experiments. JPD, APM, and LY analyzed the data. JPD, APM, and FPMV wrote the paper.

## Author Information

All data is made available at http://csbio.unc.edu/r2d2/. The authors declare no competing financial interests. Correspondence and requests for materials should be addressed to FPMV.

